# Multiple molecular and cellular properties jointly affect protein and site-specific evolutionary rates

**DOI:** 10.64898/2026.05.20.726710

**Authors:** Anshul Saini, Dinara R. Usmanova, Rydberg Supo Escalante, Dennis Vitkup

## Abstract

Protein evolutionary rates vary widely across proteins and among sites within proteins, reflecting multiple molecular, cellular, and functional constraints. While protein-level properties, such as expression and essentiality, and site-level structural and functional constraints, are known to influence evolutionary rates, how these constraints combine across scales to determine site-specific evolutionary rates remains unclear. Moreover, because many protein features are strongly correlated, it is difficult to disentangle their individual contributions to evolutionary rate variance, and unified predictive models that integrate these properties are still lacking. Here, we use neural networks to predict protein evolutionary rates across multiple scales based on multiple molecular and cellular features. At the protein level, integrating molecular and cellular descriptors explains substantial variance in evolutionary rates across proteins in multiple eukaryotic species, including nearly 50% of the variance in humans and substantial fractions of the variance in other eukaryotic species. The model also allows us to identify proteins whose evolutionary rates deviate from expectations based on their molecular and cellular properties. At the site level, we found that structural and functional features explain a comparable fraction of the variance in relative evolutionary rates. By integrating protein-level and site-level predictors, the model explains up to 37% of the variance in site-specific evolutionary rates across proteins. Our analysis demonstrates that constraints at these two scales combine largely additively, with protein-level properties setting the overall evolutionary context and site-level properties shaping variation within proteins. Together, these results provide a quantitative framework for understanding protein evolution across biological scales.

## Introduction

Understanding what determines the overall rate of protein evolution, as well as the evolution of individual sites within proteins, is a fundamental problem in evolutionary biology. Protein evolutionary rates are shaped by multiple cellular and molecular constraints, whose combined effects remain challenging to quantify (Snir et al. 2014; Zhang and Yang 2015; Biswas et al. 2018; Mawaribuchi et al. 2023; Rivas-Santisteban et al. 2025). Early theoretical work, based on neutral and nearly neutral theory, emphasized functional constraints and gene essentiality as key determinants of protein evolution (Kimura and Ohta 1974; Hurst and Smith 1999; Alvarez-Ponce et al. 2016). However, subsequent empirical studies have shown that protein expression explains the largest fraction of the variation in protein evolutionary rates, establishing the well-known expression-rate (E-R) correlation (Pál et al. 2001; Rocha and Danchin 2004). Multiple hypotheses have been proposed to explain the E-R relationship, including selection for protein stability to avoid toxic misfolding (Drummond et al. 2005; Bloom et al. 2006; Drummond and Wilke 2008) and selection for <u>f</u>unctional <u>o</u>ptimization to <u>r</u>educe the <u>c</u>osts of <u>e</u>xpression, as proposed by the FORCE mechanism (Cherry 2010; Gout et al. 2010; Usmanova et al. 2024). Consistent with the latter view, expression in neural tissues, where protein production and maintenance may be particularly costly, has been shown to impose particularly strong constraints on protein evolution in animals (Usmanova et al. 2024). Notably, several recent analyses, including ours and studies from other groups, suggest that effects related to toxic misfolding do not explain a substantial fraction of the variance in evolutionary rates across cellular proteins (Plata and Vitkup 2018; Biesiadecka et al. 2020; Usmanova et al. 2021).

In addition to expression, properties of protein interaction networks, such as connectivity and centrality, have also been shown to influence evolutionary rates (Fraser et al. 2002; Hahn and Kern 2005; Kharchenko et al. 2005; Wang et al. 2019; Mawaribuchi et al. 2023). Moreover, with the recent availability of large-scale genomic and functional datasets, these constraints can now be measured more systematically, including intolerance to loss-of-function variation from population sequencing data, such as gnomAD (Karczewski et al. 2020), as well as functional and protein– protein interaction similarity derived from Gene Ontology annotations and PPI databases (Gene Ontology Consortium 2021; Szklarczyk et al. 2023). Despite these advances, it is not fully understood how these functional and cellular properties, each imposing partially overlapping constraints on protein evolution, combine to shape protein evolutionary rates. An integrative predictive framework that brings together these intercorrelated, but potentially complementary, factors could improve our ability to explain variation in evolutionary rates and provide a more complete understanding of protein evolution. Such a model could also help identify proteins whose evolutionary behavior deviates from expectations, potentially revealing additional structural effects or function-specific selective processes.

Similar to the variance in evolutionary rates across proteins, evolutionary rates also vary substantially among residues within the same protein (Echave et al. 2016; Echave and Wilke 2017; Jayaraman et al. 2022). Sites buried in the protein core or involved in important functional regions tend to evolve more slowly than surface-exposed sites or sites with lower structural and functional importance (Franzosa and Xia 2009). While no single mechanistic model fully explains why some protein sites evolve faster than others, site-specific evolutionary rates have been shown to be strongly influenced by both structural and functional constraints (Jack et al. 2016). Among structural predictors, relative solvent accessibility (RSA) and weighted contact number (WCN) are the most important features influencing site-specific evolutionary rates across proteins and taxa (Conant and Stadler 2009; Shahmoradi et al. 2014; Yeh et al. 2014). Functional constraints also strongly influence site-specific evolutionary rates, particularly through proximity to active sites or other functionally important residues, which can generate conservation gradients across the protein (Bartlett et al. 2002; Dean et al. 2002; Abriata et al. 2015; Jack et al. 2016; Slodkowicz and Goldman 2020). Moreover, evolutionary constraint may also arise from epistatic interactions, where sites that are strongly evolutionarily coupled to other residues may experience stronger selection pressure because substitutions must remain compatible with interacting residues (Breen et al. 2012; Johnson et al. 2023). The evolutionary rate at a site is likely to depend on both global protein-level constraints and local site-level importance for maintaining biological activity. However, these two sources of constraint have been studied largely independently, and how they interact to jointly determine site-specific evolutionary rates remains poorly understood.

In this paper, we used neural networks to explain and predict protein evolutionary rates across multiple scales based on molecular, cellular, structural, and functional features. At the protein level, we modeled evolutionary rates across thousands of proteins in multiple eukaryotic species using expression, gene essentiality, protein–protein interaction (PPI) network properties, and GO-based functional similarity, with the full model explaining substantially more variance than any individual feature. At the site level, we applied a similar framework to many hundreds of enzymes to predict relative evolutionary rates within proteins using structural features, including weighted contact number (WCN) and relative solvent accessibility (RSA), as well as functional properties such as distance from the active site and epistatic interaction strength. Finally, we developed a fusion model that integrates protein-level and site-level features to predict site-specific evolutionary rates across proteins.

## Results

### Prediction of protein evolutionary rates based on their molecular and cellular properties

We first investigated whether protein evolutionary rates can be predicted from their molecular and cellular properties. To quantify how these features contribute to protein evolutionary rates, we trained neural networks using protein expression and diverse molecular and functional descriptors (Figure 1A). We first estimated empirical evolutionary rates (𝑑𝑁_𝑒𝑚𝑝_) for human proteins by calculating substitution rates, using CodeML (Yang 1997), between humans and five primate species (see Figure S1) using coding-sequence alignments obtained from the OrthoMaM database (Yang 1997; Allio et al. 2024). As predictors, we used human tissue expression profiles across 53 human tissues (Melé et al. 2015; GTEx Consortium 2020), along with gene constraint scores from the Genome Aggregation Database (gnomAD) as a measure of tolerance to likely-gene-disrupting (LGD) mutations (Karczewski et al. 2020). To capture protein function and interaction context, we incorporated GO-based functional similarity and STRING database-derived protein–protein interaction (PPI) similarity, respectively (Gene Ontology Consortium 2021; Szklarczyk et al. 2023). To reduce feature dimensionality, principal component analysis (PCA) was applied independently to expression, GO similarity, and PPI similarity matrices. For expression and GO features, PCA components explaining 99% of the variance were retained, resulting in 32 expression components and 46 GO components, whereas for the sparse PPI similarity matrices we retained a fixed number of 300 PCA components, which together explained approximately 85% of the variance (see Methods). Finally, we also used molecular descriptors for each protein, including sequence-based features such as gene length and hydrophobicity (see Methods).

**Figure 1:**
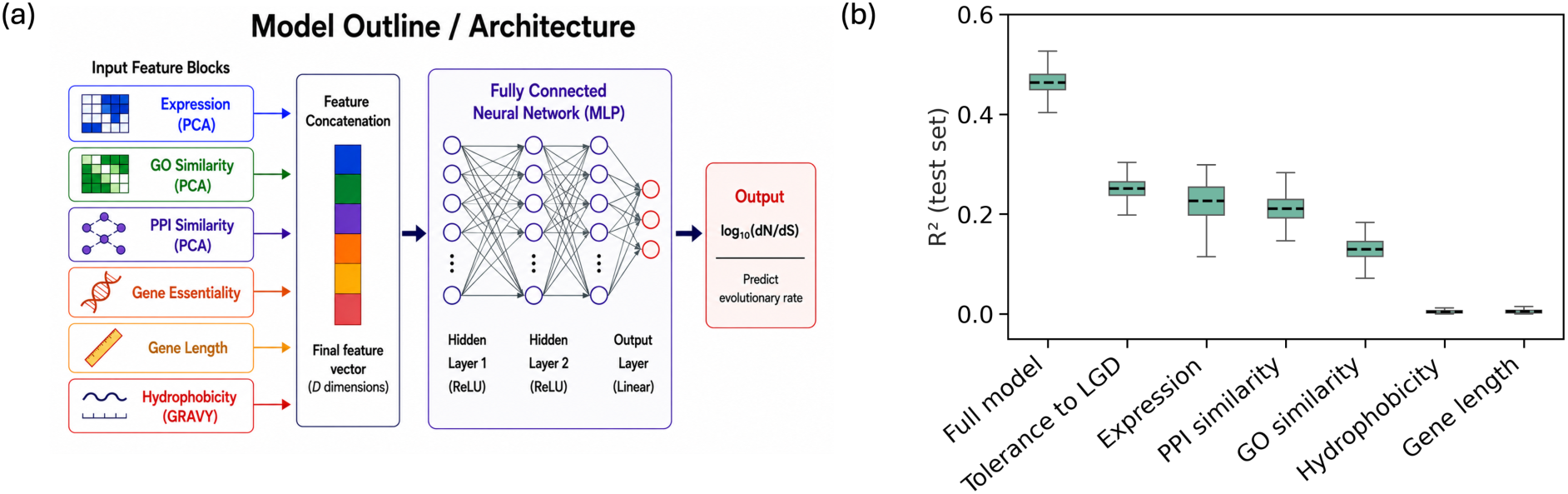
Integrated model architecture and predictive performance. (a) Schematic of the neural network used to predict protein evolutionary rates from cellular and molecular descriptors, including expression, GO-based function, PPI similarity, tolerance to LGD mutations, hydrophobicity, and gene length. The network included two hidden layers and a total of 381 input features. (b) Model performance measured as Pearson’s R^2^ on the test set for the full model and for models using individual features or feature groups. Points summarize the distribution of the R^2^ values across 500 independent runs with random splits of data into training, validation, and test sets. Boxes indicate the mean R^2^ values and standard deviations.

The features described above were used as inputs to train a neural network to predict protein evolutionary rates. Specifically, we trained a feedforward neural network with two hidden layers (containing 64 and 32 nodes) using all cellular and molecular predictors (n = 381) to estimate empirical protein evolutionary rates. To prevent data leakage, PCA components for GO-based functional similarity and PPI similarity were computed using only training genes, and GO annotations and PPI edges derived from sequence homology were excluded. Moreover, paralogous genes corresponding to genes in the test set were removed from the training and validation sets. To reduce the statistical noise in the estimation of 𝑑𝑁_𝑒𝑚𝑝_ for ultra-conserved genes, we restricted the analysis to genes with a sufficient number of substitutions (𝑁 ∗ 𝑑𝑁_𝑒𝑚𝑝_ > 0.1, where *N* is the number of sites and 𝑑𝑁_𝑒𝑚𝑝_is the substitution rate per site). Because model predictions could depend on the specific train/validation/test split, we repeated the neural network training independently 500 times, randomly reassigning genes among the training, validation, and test sets in each iteration. After filtering, an average of 13,525 human genes were retained for the analysis and split into training (80%), validation (10%), and test (10%) sets. The network was trained for up to 300 epochs with early stopping (patience = 50), using a batch size of 100, a learning rate of 10⁻³, dropout of 0.4, and weight decay of 10⁻³ (see Methods).

We found that tolerance to LGD mutations and expression were the strongest predictors of evolutionary rate, explaining 25% (𝑅^2^ = 0.25 ± 0.02) and 23% (𝑅^2^ = 0.23 ± 0.04) of the variance, respectively. These were followed by PPI similarity, which explained 21% of the variance (𝑅^2^ = 0.21 ± 0.03), and GO-based functional features, which accounted for 13% (𝑅^2^ = 0.13 ± 0.02) of the variance (Figure 1B). In contrast, molecular descriptors such as gene length and hydrophobicity explained only small fractions of variance (Figure 1B; see Table S1). When all predictors were combined, the full model explained 46% (𝑅^2^ = 0.46 ± 0.02) of the variance in protein evolutionary rates, more than double the predictive power of any individual feature (Figure 1B). Notably, brain expression, alone explained 16% of the variance, while the full model using only brain expression features still explained 45% of the variance, confirming the dominant contribution of brain expression (Drummond and Wilke 2008; Tuller et al. 2008; Usmanova et al. 2024) while also highlighting additional contributions from expression patterns across other tissues (Figure S2A).

To quantify the independent contribution of each feature, we next performed a Leave-One-Feature-Out (LOFO) analysis by retraining the network after removing one predictor at a time and measuring the reduction in the variance explained (Δ𝑅^2^). Tolerance to LGD mutations showed the largest drop (Δ𝑅^2^ = 0.09 ± 0.04), followed closely by expression (Δ𝑅^2^ = 0.07 ± 0.04), highlighting their dominant and largely independent contributions to the model prediction (Figure S3A). PPI similarity showed a moderate drop (Δ𝑅^2^ = 0.03 ± 0.05), indicating a meaningful but partially redundant contribution, while GO-based functional features showed a smaller independent effect (Δ𝑅^2^ = 0.01 ± 0.05), likely due to overlap with PPI similarity, as proteins with similar functions often share interaction partners. Interestingly, molecular features such as gene length (Δ𝑅^2^ = 0.03 ± 0.04) and hydrophobicity (Δ𝑅^2^ = 0.01 ± 0.04) showed measurable contributions despite their small standalone predictive performance, indicating that they capture orthogonal information related to molecular protein properties that is not represented by other cellular properties.

We next also evaluated feature usage by the trained network using permutation importance, in which feature values are randomized and the resulting drop in accuracy (Δ𝑅^2^) is measured. As expected, tolerance to LGD mutations (Δ𝑅^2^ = 0.25 ± 0.02) and expression (Δ𝑅^2^ = 0.19 ± 0.02) again showed the large drops (Figure S3B). Notably, PPI similarity also showed a comparable drop (Δ𝑅^2^ = 0.22 ± 0.02), despite its lower LOFO contribution. This difference indicates that while the independent signal in PPI similarity is smaller (as seen in LOFO, Figure S3A), the network still strongly relies on it when making predictions. In other words, PPI similarity provides partially shared information that is still actively used by the model in combination with other features. Similar to the LOFO analysis, GO-based functional features (Δ𝑅^2^ = 0.08 ± 0.03) and molecular features such as gene length (Δ𝑅^2^ = 0.08 ± 0.02) and hydrophobicity (Δ𝑅^2^ = 0.031 ± 0.007) showed substantially weaker effects (Figure S3B).

Overall, these results suggest that protein evolutionary rates are shaped by multiple molecular and cellular properties, and that complementary biological features substantially improve the prediction of evolutionary rates. While features such as expression and tolerance to LGD mutations provide strong independent signals, PPI similarity and GO descriptors capture overlapping biological information that reflects shared functional relationships among proteins. In contrast, molecular descriptors generally contribute weaker but complementary information.

### Predictions of evolutionary rates for individual proteins

The predictive model incorporates multiple functional and molecular descriptors to estimate evolutionary rates for each protein. Therefore, it can also be used to identify proteins whose evolutionary behavior is not well explained by the features included in the model. Proteins that deviate, in terms of their evolutionary rate, from others with similar cellular and molecular descriptors are expected to be poorly predicted, suggesting additional evolutionary constraints or processes not captured by the overall model. To explore these mispredictions, we computed the evolutionary rate prediction error (Δ𝑑𝑁 = 𝑙𝑜𝑔_10_(𝑑𝑁_𝑝𝑟𝑒_/𝑑𝑁_𝑒𝑚𝑝_;) for each protein. This score was averaged across 500 independent model-training runs to estimate 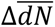, with each run based on a random reassignment of proteins into training, validation, and test sets (Figure 2A, B).

**Figure 2:**
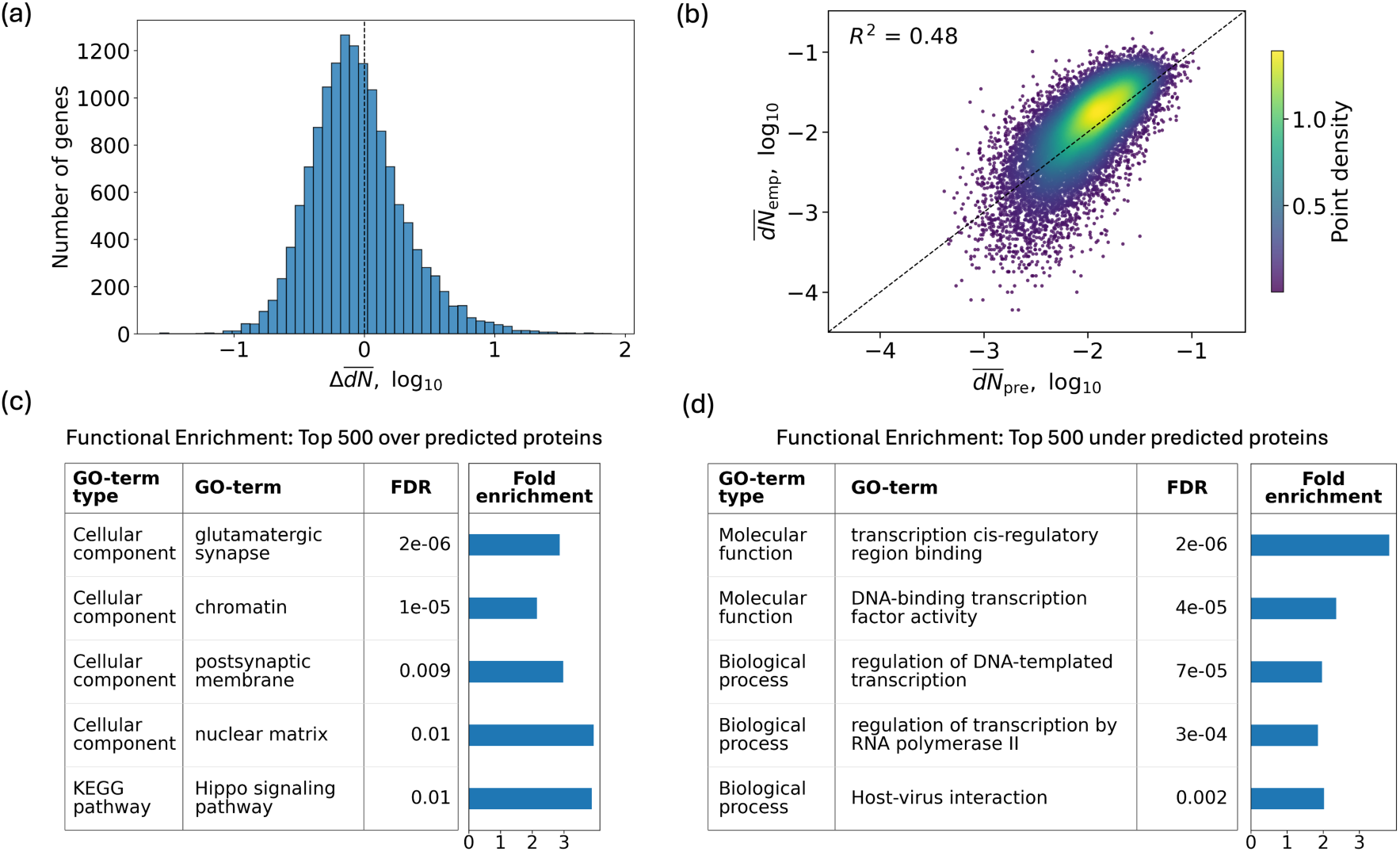
Gene-level prediction accuracy and functional characterization of outlier genes. (a) Histogram of the average prediction error, 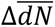, in nonsynonymous evolutionary rates aggregated across 500 runs. For each run, the prediction error was defined as Δ𝑑𝑁 = log_10_(𝑑𝑁_𝑝𝑟𝑒_/ 𝑑𝑁_𝑒𝑚𝑝_;, where 𝑑𝑁_𝑝𝑟𝑒_ is the model predicted substitution rate at nonsynonymous sites and 𝑑𝑁_𝑒𝑚𝑝_is the empirical substitution rate estimated using a star-shaped phylogeny across primate species (see Methods).(b) Density scatter plot comparing predicted (𝑑𝑁_𝑝𝑟𝑒_) and empirical evolutionary rates (𝑑𝑁_𝑒𝑚𝑝_) across all genes, showing mean values across runs. The dashed line indicates the identity line. GO enrichment analysis of the (c) top 500 overpredicted (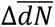> 0) and (d) underpredicted (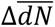< 0) genes, ranked by mean prediction error. Functional enrichment was calculated using DAVID, and only GO terms with Benjamini–Hochberg adjusted p-values FDR < 0.02 and annotated with a minimum of 50 and a maximum of 2000 genes are shown to avoid overly general or highly specific terms.

To examine the functional bias of outlier proteins, we then selected the top 500 overpredicted and underpredicted proteins from the model and performed functional enrichment analysis using the DAVID functional annotation tool (Sherman et al. 2022). We found that the overpredicted proteins, i.e., proteins evolving slower than the model predicted rate (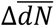> 0), are enriched for several neuronal functions, including glutamatergic synapse and postsynaptic membrane proteins. This suggests that some neuronal proteins experience stronger functional constraints than are generally captured by the model (Figure 2C, see Table S2). Overpredicted proteins were also enriched in the Hippo signaling pathway, a highly conserved pathway involved in regulating organ growth through the control of cell proliferation and apoptosis. In contrast, underpredicted proteins, i.e., proteins evolving faster than the model predicted rate(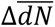< 0), were enriched for transcriptional regulators, DNA-binding proteins, and host–virus interaction pathways (Figure 2D, see Table S3). This likely reflects shifting adaptive selection pressures and regulatory constraints that are not fully captured by typical cellular predictors such as expression, PPI similarity, and GO functional descriptors.

### Prediction of protein evolutionary rate across multiple eukaryote species

To test the generality of our prediction framework, we extended the model to multiple eukaryotic species to examine whether protein evolutionary rates can be inferred from molecular and cellular descriptors across diverse taxa. Specifically, in addition to *Homo sapiens*, we applied the approach to *Mus musculus*, *Drosophila melanogaster*, and *Arabidopsis thaliana*. For each of these species, the empirical evolutionary rates, 𝑑𝑁_𝑒𝑚𝑝_were calculated using CodeML based on sets of closely related species (Figure S1, see Methods). As predictors, we used species-specific tissue expression profiles, gene essentiality measures, PPI similarity, and GO-based functional similarity, along with molecular descriptors, such as protein hydrophobicity and length (see Methods). For expression datasets, we used expression profiles across 13 tissues for *Mus musculus* (Söllner et al. 2017), 18 tissues for *Drosophila melanogaster* (Leader et al. 2018), and 79 tissues for *Arabidopsis thaliana* (Klepikova et al. 2016). Gene essentiality descriptors were derived from phenotype-based annotations using species-specific resources: genotype–phenotype and viability data from *FlyBase* for *Drosophila melanogaster* (Larkin et al. 2021), mutant phenotype and developmental lethality datasets for *Arabidopsis thaliana* (Berardini et al. 2015), and viability-based knockout phenotype annotations for *Mus musculus* (Dickinson et al. 2016) (see Methods). Similar to the human analysis, PCA-based dimensionality reduction was applied to expression, GO similarity, and PPI similarity features, retaining components explaining 99% of the variance for expression and GO features and 300 components for sparse PPI similarity matrices (see Methods). Molecular descriptors for each protein, including protein length and protein hydrophobicity, were inferred from amino acid sequences (see Methods).

The full model explained a substantial fraction of protein evolutionary rate variation in all species considered, with 38% of the variance explained (𝑅^2^ = 0.38 ± 0.03) in *Mus musculus*, 33% (𝑅^2^ = 0.33 ± 0.03) in *Drosophila melanogaster*, and 39% (𝑅^2^ = 0.39 ± 0.02) in *Arabidopsis thaliana* (Figure 3). The relative contributions of predictors were broadly consistent across the species, with expression emerging as the strongest determinant of evolutionary rate, consistent with previous studies (Pál et al. 2001; Drummond and Wilke 2008; Usmanova et al. 2024). Specifically, expression explained 20%, 24%, and 30% of the protein evolutionary rate variance in *Mus musculus, Drosophila melanogaster*, and *Arabidopsis thaliana,* respectively (Figure 3). Moreover, the largest tissue-specific contributions were derived from brain expression in *Mus musculus* and *Drosophila melanogaster*, and from germinating seedling expression in *Arabidopsis thaliana*, with these tissues individually explaining 18%, 18%, and 23% of the variance, respectively. The full model using expression features from only these tissues was still able to explain comparable amounts of variance, achieving 𝑅^2^ values of 37%, 29%, and 37% for *Mus musculus*, *Drosophila melanogaster*, and *Arabidopsis thaliana*, respectively (Figure S2). Expression was followed by PPI similarity and GO-based functional features, whereas gene essentiality and molecular descriptors were comparatively weaker predictors. Notably, gene essentiality based on viability data in *Drosophila melanogaster* (𝑅^2^ = 0.07 ± 0.01) showed stronger predictive power than in *Mus musculus* (𝑅^2^ = 0.023 ± 0.008) and *Arabidopsis thaliana* (𝑅^2^ = 0.004 ± 0.003), possibly reflecting differences in annotation quality, phenotype coverage, and the biological specificity of essentiality measures across species.

**Figure 3:**
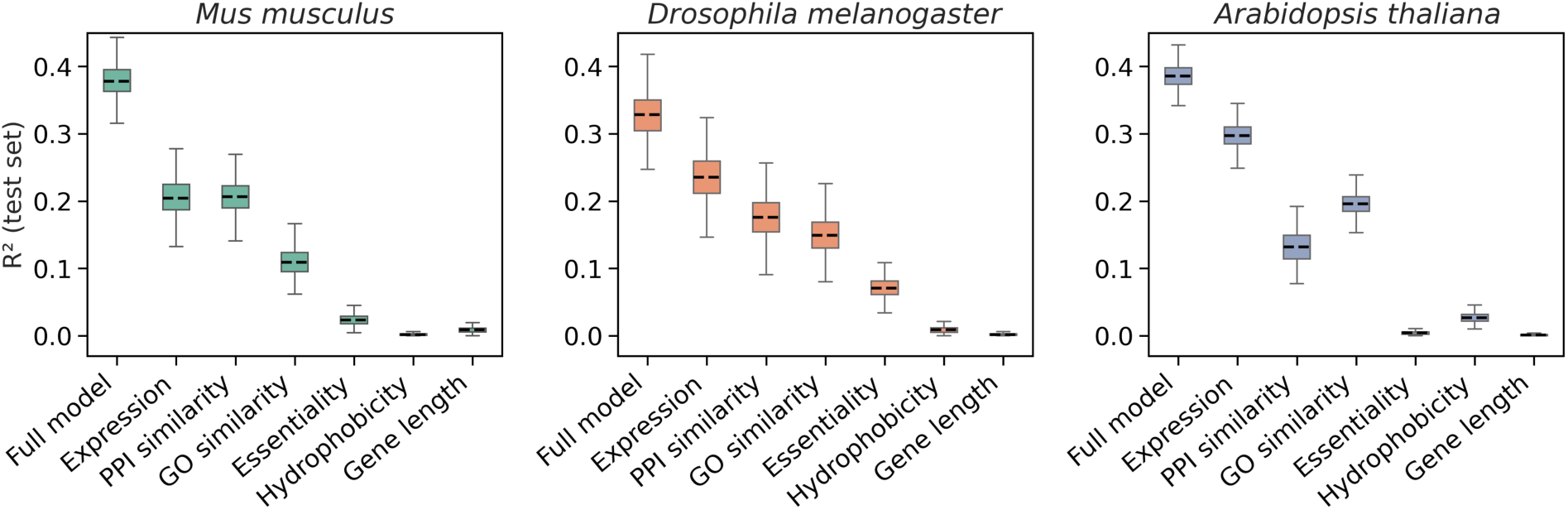
Integrated model performance across diverse eukaryotic species. Predictive performance measured as test-set R^2^ of the full model and individual feature groups across representative eukaryotic lineages: *Mus musculus* (mammals), *Drosophila melanogaster* (invertebrates), and *Arabidopsis thaliana* (plants). Empirical evolutionary rates for each species were estimated using star-shaped phylogenies (see Methods). For each model and each feature group, points summarize the distribution of R^2^ values across 500 independent runs with random split of data into training, validation, and test gene sets. Boxes in the panels indicate the mean R^2^ values and standard deviations.

We further performed LOFO and permutation analyses to assess the importance and usage of each predictive feature across species. Consistently, expression showed the largest drop in explanatory power (Δ𝑅^2^) in the LOFO analysis with Δ𝑅^2^ = 0.10 ± 0.05 in *Mus musculus,* 0.10 ± 0.05 𝑖𝑛 *Drosophila melanogaster*, and 0.12 ± 0.02 in *Arabidopsis thaliana* (Figure S4). Similarly, permutation analysis, which reflects feature usage by the trained model, showed that expression was the most utilized feature across species. In particular, for *Arabidopsis thaliana*, permuting the expression block led to a very strong reduction of predictive power, indicating that the model relies heavily on expression in this species (Figure S5). Overall, these results demonstrate that protein evolutionary rates are governed by broadly similar molecular, cellular, and functional features across species, and that integrating these features enables substantial improvements in the fraction of evolutionary rate variance explained in diverse eukaryotes.

### Prediction of site-specific evolutionary rates across proteins using their molecular and cellular properties

Our analyses demonstrated that protein evolutionary rates can be predicted using molecular and cellular properties of proteins. However, substantial variation in evolutionary rates also exists within proteins, where different sites experience distinct local constraints. Previous work has shown that site-specific evolutionary rates are influenced by structural, functional, and epistatic constraints (Breen et al. 2012; Jack et al. 2016), but how these local constraints combine with broader protein-level properties to influence site-specific evolutionary rates remains unclear. To address this, we first trained a neural network to predict relative evolutionary rates across sites using structural, functional, and epistatic predictors (Figure S6A). We then implemented a fusion neural network that integrates protein-level and site-level predictors to model site-specific evolutionary rates across multiple proteins (Figure 4A).

**Figure 4:**
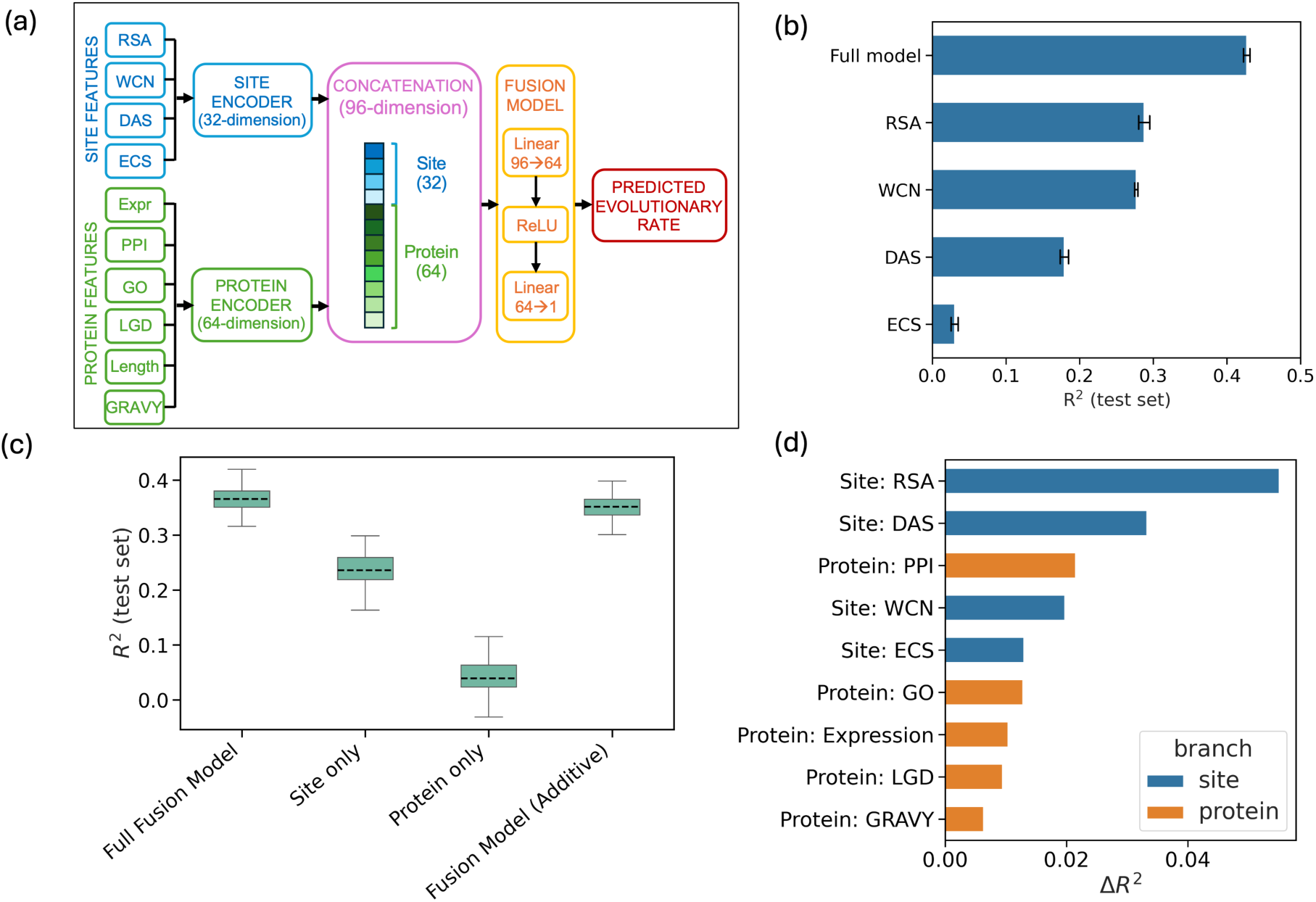
Multiscale modeling of site-specific evolutionary rates across proteins. (a) Schematic of the full fusion neural network architecture. Site-level features included relative solvent accessibility (RSA), weighted contact number (WCN), distance from the active site (DAS), and epistatic coupling strength (ECS) and protein-level features (Expression (Expr), GO, PPI, Tolerance to LGD mutations (LGD), hydrophobicity measured using the GRAVY score and Protein length (Length)) were encoded separately using a site-specific encoder (32 dimensions) and a protein-specific encoder (64 dimensions). The encoded representations were concatenated and passed through a fusion network to predict site-specific evolutionary rates across proteins. (b) Performance of a standalone site-level neural network (Figure S6A) trained using only structural and epistatic features to predict relative evolutionary rates within proteins. The test-set 𝑅^2^ is shown for the full site model and for individual features (RSA, WCN, DAS, ECS). (c) Predictive performance of the full fusion model compared with reduced models (site-only, protein-only, and additive fusion models). Boxplots show the distribution of R^2^ values across 100 independent runs with genes randomly split into training, validation, and test gene sets. (d) Feature importance in the fusion model based on leave-one-feature-out (LOFO) analysis, showing the contribution of site-level (blue) and protein-level (orange) features to the model performance, measured as decrease in *R^2^* (Δ𝑅^2^) after the corresponding feature removal.

First, we trained a neural network to predict the relative evolutionary rates across protein sites using their structural and functional properties (Figure S6A). We focused on enzymes because their catalytic residues and active sites are comparatively well annotated, allowing us to quantify the contribution of evolutionary constraints related to active site proximity. To estimate relative evolutionary rates across sites, we selected 100 eukaryotic species and retrieved their complete proteome datasets from UniProt (UniProt Consortium 2025). For each conserved metabolic enzyme, orthologous sequences across species were identified using DIAMOND sequence similarity searches (Buchfink et al. 2021). After alignment and structural filtering, we estimated relative evolutionary rates using Rate4Site (Pupko et al. 2002) for the remaining 51,573 sites across 216 proteins, where site rates were estimated as the evolutionary rate of a site relative to the mean rate of the corresponding protein (see Methods). Following previous work, we used structural features related to protein packing density, WCN and RSA, as predictors for each protein site (Jack et al. 2016). To capture the effect of catalytic constraints, we also calculated the distance from the active site for each enzyme using available PDB structures (Burley et al. 2019). Epistatic effects were incorporated by estimating the mean epistatic coupling strength (ECS) of each site with all other protein sites using data obtained from EVmutation (see Methods) (Hopf et al. 2017). The site-level features and their corresponding relative evolutionary rates from all proteins were then concatenated and used to train a neural network. For the neural network, we used a three-layer architecture with 128, 64, and 32 nodes. For training, we used up to 300 epochs with early stopping (patience = 50), a batch size of 100, a learning rate of 10⁻³, weight decay of 10⁻³, and a dropout rate of 0.1 (see Methods).

Consistent with previous results (Jack et al. 2016), we found that RSA explained 29% (𝑅^2^ = 0.288 ± 0.008), the highest fraction of variance in evolutionary rates across sites, closely followed by WCN, which explained 28% (𝑅^2^ = 0.277 ± 0.002), while distance from the active site (DAS) explained a slightly smaller fraction 18% (𝑅^2^ = 0.179 ± 0.006) of the variance on its own (Figure 4B). Epistatic coupling strength (ECS) by itself explained only a small fraction, 3% (𝑅^2^ = 0.030 ± 0.005), of the evolutionary rate variance. The full model incorporating all these features explained 43% of the variance (𝑅^2^ = 0.427 ± 0.004) in site-specific evolutionary rates, indicating that these predictors provide complementary information when combined. Furthermore, DAS showed the largest LOFO contribution (Δ𝑅^2^∼ 0.07), indicating that it provides a strong functional signal not captured by packing-based properties such as WCN and RSA (Figure S6B).

Similarly, epistatic interactions also contributed unique information to the prediction. In contrast, WCN showed the lowest LOFO contribution (Δ𝑅^2^∼0.02), suggesting that much of its information overlaps with RSA. Consistent with these results, permutation importance analysis identified RSA and DAS as the features most strongly used by the network to predict site-specific evolutionary rates (Figure S6C).

Notably, relative evolutionary rates across sites within a protein do not capture how site evolution is affected by cellular and functional properties of the corresponding protein. To incorporate both protein-level and site-level effects, we next estimated site-specific evolutionary rates across multiple proteins. To that end, we randomly selected 50 sites from each of 1083 enzymes across 100 species, concatenated them into a single alignment, and estimated evolutionary rates across all these sites from multiple proteins simultaneously using IQ-TREE. In this way, the estimated rate at each site reflects both the overall conservation of the protein and the local constraints acting at each site. To predict site-specific evolutionary rates across proteins, we implemented a fusion neural network with two components: a protein encoder that learns from protein-level features and a site encoder that captures within-protein variation using site-level predictors (Figure 4A). A fusion head was then used to integrate the representations from both encoders and was trained on the combined dataset to predict site-specific evolutionary rates across proteins. The protein encoder was trained using the protein-level features (Figure 1A) and CodeML-based protein evolutionary rates, whereas the site encoder was trained using the site-level features and relative site-specific rates estimated using IQ-TREE. The fusion head was then trained using site-specific evolutionary rates calculated using the concatenated alignments of sites from multiple proteins; these site-specific rates reflect both protein-level and site-level effects (see Methods). All neural networks were trained using the Adam optimizer with mean-squared-error loss. The protein and site encoders were trained for up to 300 epochs with a learning rate of 10⁻³, weight decay of 10⁻³, dropout of 0.2, batch sizes of 256 and 1024 for protein and site models, respectively, and early stopping with a patience of 50 epochs based on validation loss. The complete training and evaluation procedure was repeated across 100 independent random train/validation/test gene splits.

The developed fusion model explained 37% (𝑅^2^ = 0.37 ± 0.02) of the variance in site-specific evolutionary rates across proteins (Figure 4C). Using LOFO analysis, we found that RSA displayed the largest drop in variance explained (Δ𝑅^2^∼ 0.06), followed by distance from the active site (Δ𝑅^2^∼ 0.03) and PPI similarity (Δ𝑅^2^∼ 0.02), indicating that both local structural constraints and protein-level functional constraints contributed to the prediction (Figure 4D). To assess the relative contribution of protein-level and site-level information, we performed an ablation analysis in which features associated with each encoder were used separately to predict the output. Site-level features alone explained 25% of the variance, whereas protein-level features alone explained 4%, indicating a substantially stronger contribution from site-specific information for this prediction task. This is consistent with ANOVA-based variance decomposition (Searle et al. 2006) (see Methods), which showed that most of the variance in evolutionary rates lies within proteins across sites (79%) rather than between proteins (21%).

We also tested whether interactions between protein-level and site-level predictors are strongly non-linear. For this, we trained a reduced fusion model with a single-layer fusion head, allowing only additive integration of the protein-level and site-level effects. The variance explained by the additive model (𝑅^2^ = 0.35 ± 0.02) was only slightly lower than that of the full nonlinear model with three layers, suggesting that the contributions from protein-level and site-level predictors can be captured through additive effects (Figure 4C). Overall, these results indicate that both site-specific structural and functional constraints and protein-level properties contribute to evolutionary rates, with site-level features shaping variation among residues within proteins, while protein-level properties such as expression and functional descriptors provide the broader evolutionary context in which those local constraints act.

## Discussion

Protein evolutionary rates are shaped by multiple molecular, structural, and cellular constraints acting across different biological scales (Zhang and Yang 2015; Echave et al. 2016; Usmanova et al. 2024). Rather than being determined by a single dominant factor, our results suggest that evolutionary rates emerge from the combined effects of multiple, partially overlapping molecular, cellular, and functional constraints. While individual predictors, such as expression, tolerance to LGD mutations, and PPI similarity, explain substantial fractions of evolutionary rate variance, no single feature accounts for the majority of rate variation. However, by utilizing a neural network to integrate multiple predictors, we were able to explain nearly half of the variance in human protein evolutionary rates (∼46%). The doubling of predictive power in the full model compared to any standalone feature, suggests that the large variation in protein evolutionary rates likely reflects the collective influence of multiple constraints associated with protein function, interaction context, and cellular properties. The strong contribution of expression-based predictors is consistent with the FORCE mechanism, which links high protein expression to stronger evolutionary conservation through selection on functional efficiency (Cherry 2010; Gout et al. 2010; Usmanova et al. 2024). Moreover, within this framework, the added predictive value of functional and interaction-related features may further reflect variation the strength of selection for functional optimization across proteins.

An important aspect of the integrated predictive framework is that it allows us to identify proteins whose evolutionary rates deviate from expectations based on their molecular and cellular properties. Specifically, proteins associated with neuronal function were consistently overpredicted by the model, suggesting that they frequently exhibit stronger evolutionary conservation than expected from the cellular and molecular features included in the model. In contrast, genes involved in transcriptional regulation and immunity tend to be underpredicted, consistent with the fact that these proteins often experience adaptive or lineage-specific changes in selective pressures not captured by steady-state cellular descriptors (Rockman et al. 2003; Carroll 2005; Barreiro and Quintana-Murci 2010). These deviations suggest that integrative evolutionary rate models can help to identify biological processes and functions associated with unusual evolutionary dynamics beyond general cellular constraints.

At the protein site level, structural constraints explain a large fraction of the variance in relative evolutionary rates across protein sites. In particular, WCN and RSA act as strong predictors of evolutionary rate, consistent with the idea that local packing density and solvent exposure strongly influence the fitness effects of mutations. However, structural properties alone do not fully explain site-specific evolutionary rates. Functional constraints provide an additional layer of selection, as reflected by the contribution of distance to active sites and epistatic interaction strength. Although distance to the active site is a weaker predictor on its own, it captures constraints associated with catalytic importance that extend beyond the active site itself (Jack et al. 2016). Similarly, epistatic interactions contribute information beyond structural features, suggesting that evolutionary rates are also influenced by compatibility constraints among interacting residues arising from coevolution. Importantly, our results show that evolutionary rates at individual sites are influenced by both protein-level and site-level constraints. A key finding is that these two levels combine largely additively to determine evolutionary rates across sites. This suggests that much of the variation in evolutionary rates can be decomposed into a global protein-level component and a local site-specific component.

Applying the predictive framework to different species also highlights the importance of data quality and functional annotation. In humans, gene essentiality, as measured through intolerance to LGD mutations, is a strong predictor of evolutionary rate, whereas in other species its contribution is substantially weaker, likely due to limitations in available functional genomic and knockout datasets. As genomic and functional annotations improve across species, integrative frameworks for predicting evolutionary rates should capture a larger fraction of the evolutionary rate variance and better resolve the underlying constraints shaping protein evolution. Finally, although the current analysis focused on multicellular eukaryotes, the developed framework could be extended to unicellular organisms, where additional evolutionary processes, such as codon adaptation, horizontal gene transfer, and operon-level selection, may play important roles in shaping protein evolutionary rates (Koonin et al. 2001; Rocha 2004; Price et al. 2006). Incorporating these factors could provide a broader and more unified understanding of the factors affecting protein evolutionary rate across all domains of life.

## Methods

### Estimation of protein evolutionary rate

To estimate protein evolutionary rates for mammalian species, we used one-to-one orthologous coding sequence alignments from the OrthoMaM database (Allio et al. 2024), which provides codon alignments across 190 mammalian species. For Human (*Homo sapiens*), we selected five progressively diverging primate species: *Gorilla gorilla*, *Pongo abelii*, *Macaca mulatta*, *Aotus nancymaae*, and *Microcebus murinus* (Figure S1). For Mouse (*Mus musculus*), we selected *Mus caroli, Mus pahari, Grammomys surdaster, Rattus norvegicus, and Acomys russatus* (Figure S1). For each focal species, pairwise nonsynonymous divergence (*dN*) was estimated between the focal species and each of the five selected related species using Codeml (runmode = -2) from the PAML package (Yang 1997). Protein evolutionary rates (𝑑𝑁_𝑒𝑚𝑝_) were then estimated for each protein as the mean pairwise *dN* across all five species comparisons. For *Drosophila melanogaster* and *Arabidopsis thaliana*, protein evolutionary rates estimates were obtained from a recent study (Usmanova et al. 2024), where 𝑑𝑁_𝑒𝑚𝑝_ for *Drosophila melanogaster* were estimated using pairwise *dN* from *Drosophila yakuba*, and 𝑑𝑁_𝑒𝑚𝑝_for *Arabidopsis thaliana* were estimated using pairwise *dN* from *Arabidopsis halleri, Brassica oleracea, Glycine max, Solanum lycopersicum,* and *Helianthus annuus*.

### Protein evolution rate predictor data

Human expression profiles across 53 tissues were obtained from the GTEx portal (GTEx Consortium 2020), mouse expression data for 13 tissues from ArrayExpress (Söllner et al. 2017), and *Drosophila melanogaster* larval expression data for 18 tissues from FlyAtlas2 (Leader et al. 2018). *Arabidopsis thaliana* expression profiles across 79 tissues were obtained from a previously reported RNA-seq dataset (Klepikova et al. 2016). Expression values were averaged across replicates or samples for each tissue. Functional and protein–protein interaction (PPI) data were derived from Gene Ontology (GO) annotations (Gene Ontology Consortium 2021) and the STRING database (Szklarczyk et al. 2023), respectively. To minimize sequence-based information leakage, GO annotations inferred from sequence similarity (ISS, ISO, ISA, ISM, IBA, IBD, IKR, IRD, IEA, and RCA) were excluded. Similarly, for PPI, we only retained high-confidence interactions (score >700) and removed interactions with non-zero homology scores to further avoid sequence-based leakage.

Gene essentiality scores for humans were obtained from the gnomAD constraint metrics database (v4.1). We used the upper bound of the loss-of-function observed/expected confidence interval (LOEUF) for canonical transcripts as a measure of intolerance to loss-of-function mutations, where lower LOEUF values indicate stronger functional constraint. For other species, gene essentiality measures were derived from phenotype-based annotations using species-specific resources: genotype–phenotype and viability data from *FlyBase* for *Drosophila melanogaster* (Larkin et al. 2021), mutant phenotype and developmental lethality annotations for *Arabidopsis thaliana* (Berardini et al. 2015), and viability-based knockout phenotype annotations for *Mus musculus* (Dickinson et al. 2016). Protein sequence-based descriptors were calculated from reference proteomes obtained from Ensembl. For each gene, the canonical protein isoform, or the longest isoform when canonical annotation was unavailable, was selected. Hydrophobicity was estimated using the GRAVY score calculated with the ProteinAnalysis module in BioPython, and the fraction of charged amino acids was calculated as the proportion of D, E, K, and R amino acids in the sequence.

### Neural Network for protein evolutionary rate prediction

We trained species-specific fully connected feedforward neural networks with two hidden layers (64 and 32 nodes) to predict protein evolutionary rates (𝑑𝑁_𝑒𝑚𝑝_) from cellular and molecular features. Predictors included tissue-specific expression profiles, GO-based functional similarity, protein–protein interaction (PPI) similarity, gene essentiality scores, and sequence-based molecular descriptors such as protein length and hydrophobicity. Species-specific tissue expression datasets included 53 tissues for *Homo sapiens*, 13 for *Mus musculus*, 18 for *Drosophila melanogaster*, and 79 for *Arabidopsis thaliana*. To reduce feature dimensionality, principal component analysis (PCA) was applied independently to the expression, GO similarity, and PPI similarity matrices for each species. GO-based functional similarity and PPI similarity matrices were constructed using cosine similarity between gene annotation and interaction profiles, respectively. For expression and GO features, components explaining 99% of the variance were retained. This resulted in 32 expression PCs and 46 GO PCs for *Homo sapiens*, 11 expression PCs and 32 GO PCs for *Mus musculus*, 16 expression PCs and 33 GO PCs for *Drosophila melanogaster*, and 67 expression PCs and 18 GO PCs for *Arabidopsis thaliana*. Because the PPI similarity matrices were highly sparse and disconnected, variance-based dimensionality reduction produced unstable high-dimensional representations. Therefore, we retained a fixed number of 300 PCA components for PPI features across all species, which together explained approximately 85% of the variance in the PPI similarity matrices. Gene essentiality scores and sequence-based molecular descriptors were included as additional scalar predictors.

To prevent data leakage, PCA components were calculated using only training genes, and paralogous genes overlapping between the training and validation/test sets were excluded. In total, the model used 381 predictors for *Homo sapiens*, 346 for *Mus musculus*, 352 for *Drosophila melanogaster*, and 388 for *Arabidopsis thaliana*. To reduce statistical noise in 𝑑𝑁_𝑒𝑚𝑝_estimates, we retained only proteins with more than 200 synonymous and nonsynonymous sites. Because primates accumulate fewer substitutions due to their longer generation times, additional filtering was applied to remove highly conserved genes with insufficient substitutions (𝑁 ∗ 𝑑𝑁_𝑒𝑚𝑝_ > 0.1, where *N* is the number of nonsynonymous sites and 𝑑𝑁_𝑒𝑚𝑝_ is the nonsynonymous substitution rate per site). Because the set of retained genes depended on the train/validation/test split following paralog removal, the number of genes varied slightly across runs. On average, the final datasets contained 13,525 genes for *Homo sapiens*, 13,725 for *Mus musculus*, 12,051 for *Drosophila melanogaster*, and 16,642 for *Arabidopsis thaliana*. Genes were randomly divided into training, validation, and test sets using an 80/10/10 split. To account for variability arising from random dataset partitioning, the neural network training was independently repeated 500 times with new random gene splits in each iteration. Models were trained for up to 300 epochs using a learning rate 10^-3^, dropout 0.4, and weight decay 10^-3^. Early stopping based on validation loss was applied with a patience of 50 epochs. To further examine the contribution of dominant tissues driving the expression–evolutionary rate (E–R) relationship, we additionally repeated the analysis using expression data from a single tissue. These single-tissue models used brain expression for *Homo sapiens, Mus musculus* and *Drosophila melanogaster*, and germinating seed expression for *Arabidopsis thaliana*.

### Individual protein prediction analysis

To identify the mis-predicted proteins, for each protein, we estimated the average misprediction error (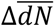) by the model. For that, we first calculate the misprediction errors for every protein as 𝛥d𝑁 = log 10(𝑑𝑁_𝑝𝑟𝑒_/𝑑𝑁_𝑒𝑚𝑝_;. Because misprediction of a protein in the test set can depend on the proteins selected in the training set. We averaged this score across 500 independent runs with randomly splitting genes into training/validation/test split to estimate the average misprediction error (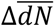). Genes with positive 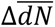 values were classified as overpredicted, whereas genes with negative values were classified as underpredicted. To characterize functional biases among mis-predicted proteins, we performed DAVID functional enrichment analysis (Sherman et al. 2022) separately for the top 500 overpredicted and underpredicted genes. DAVID enrichment results were filtered to retain terms with Benjamini-adjusted FDR < 0.02 and population counts between 50 and 2000 genes to exclude overly broad or sparsely represented functional categories. We additionally excluded sequence-feature, ligand-binding, and disease-related annotations to focus on broader biological and functional processes.

For visualization purposes (Figure 2), representative enriched terms were selected from the filtered DAVID results. For overpredicted genes, selected terms included glutamatergic synapse, synapse, postsynaptic membrane, chromatin, and Hippo signaling pathway. For underpredicted genes, selected terms included DNA-binding transcription factor activity, transcription cis-regulatory region binding, host–virus interaction, regulation of DNA-templated transcription, and regulation of transcription by RNA polymerase II. Fold-enrichment values were plotted for the selected terms, and DAVID annotation categories were grouped into broader functional classes such as molecular function, biological process, cellular component, and KEGG pathway. All enriched terms passing the filtering criteria were provided in Supplementary Tables (Table S2, S3).

### Prediction of relative evolutionary rate across sites within proteins

We selected metabolic enzymes (n = 1,990) for which experimentally annotated active-site positions were available, allowing us to calculate the structural distance of each residue from the nearest active site. To construct multi-species alignments, we downloaded eukaryotic reference proteomes from UniProt (UniProt Consortium 2025) and randomly selected 100 non-human eukaryotic species. Human protein sequences derived from AlphaFold structures were used as queries for DIAMOND BLAST (Buchfink et al. 2021) searches against each selected proteome to identify homologous proteins. Protein alignment containing homologs from fewer than 80 species were excluded, resulting in a final dataset of 574 proteins. For each protein alignment, homologous sequences across species were aligned using Clustal Omega (Sievers and Higgins 2018). Relative evolutionary rates across sites were then estimated for each protein alignment using Rate4Site (Pupko et al. 2002) under a maximum-likelihood framework.

Structural features including relative solvent accessibility (RSA) and weighted contact number (WCN) were calculated from AlphaFold structures using a previously developed structural analysis pipeline (Jack et al. 2016). To capture catalytic constraints, we estimated the distance of each residue from experimentally annotated active-site residues using protein structures and active-site annotations available from the PDB database (Burley et al. 2019). Epistatic constraints were incorporated using evolutionary coupling data obtained from EVmutation (Hopf et al. 2017). For each residue, we calculated the total epistatic coupling strength (ECS) as the sum of the absolute coupling scores |cn| between that residue and all other residues within the same Pfam domain.

We trained a feedforward neural network with three hidden layers (128, 64, and 32 nodes) to predict relative site-specific evolutionary rates within proteins. For each residue, input features included WCN, RSA, DAS, and ECS, while relative evolutionary rates estimated using Rate4Site were used as the target variable. To ensure reliable structural and evolutionary estimates, we retained only residues with high AlphaFold confidence scores (pLDDT > 80) and alignment positions containing no more than five gaps across the multi-species alignment. After applying structural and alignment quality filters, data from all proteins were concatenated into a single dataset consisting of approximately 51573 sites across 216 proteins. The dataset was randomly partitioned into training (80%), validation (10%), and test (10%) sets. The neural network was trained for up to 300 epochs using a dropout rate of 0.1, batch size of 100, learning rate 10^-3^, and weight decay 10^-3^.

### Prediction of site-specific evolutionary rates across the proteins

To predict site-specific evolutionary rates across proteins, we implemented a multimodal neural network that integrates protein-level and site-level predictors. The model consisted of a protein encoder, a site encoder, and a fusion head. The protein encoder was pretrained to predict protein-level evolutionary rates from expression, GO-based functional components, PPI components, gene essentiality, protein length, and hydrophobicity. To avoid overparameterization in the fusion model, high-dimensional protein-level predictors were compressed using PCA, retaining 30 principal components each for expression, GO similarity, and PPI similarity features. The site encoder was pretrained to predict within-protein relative evolutionary rates derived from Rate4Site using RSA, WCN, DAS, and ECS as input features. The protein encoder used two hidden layers with 128 and 64 nodes, whereas the site encoder used two hidden layers with 64 and 32 nodes. Both encoders used ReLU activations and dropout regularization.

To train the fusion model, we constructed a combined dataset containing both inter-protein and intra-protein variation in evolutionary rates. To avoid overrepresentation of large proteins and maintain computational tractability, we randomly sampled 50 sites from each enzyme alignment spanning 100 species, retaining only sites with high-confidence AlphaFold predictions (pLDDT ≥ 80) and fewer than 20 gaps across the alignment. The sampled sites were concatenated into a single meta-alignment comprising 54,048 sites derived from 1,083 proteins. A phylogenetic tree for the sampled species was inferred from the meta-alignment using IQ-TREE under the LG+R10 model (Minh et al. 2020). Using this tree as a fixed topology, site-wise evolutionary rates across the concatenated alignment were estimated using IQ-TREE with the -wsr option under the LG+G20 model. For each sampled site, protein-level and site-level predictors were combined to predict the corresponding IQ-TREE site-wise evolutionary rate. To quantify the relative contributions of protein-level and site-level variation, we performed a one-way ANOVA-based variance decomposition using protein identity as the grouping factor. Site-wise evolutionary rates estimated by IQ-TREE were partitioned into between-protein and within-protein components. This analysis was used to estimate the fraction of evolutionary-rate variation attributable to differences between proteins versus variation among sites within proteins.

The fusion model was initialized using the pretrained protein and site encoders. During fusion training, encoder weights were initially frozen and only the fusion head was trained for up to 60 epochs. The model was subsequently fine-tuned by unfreezing the encoder linear layers and jointly training both encoders and the fusion head for up to 60 additional epochs. To avoid information leakage between homologous sites from the same protein, proteins rather than individual sites were partitioned into training (80%), validation (10%), and test (10%) sets. Early stopping based on validation loss was applied with a patience of 12 epochs during fusion-head training and 10 epochs during fine-tuning. The model was trained using a batch size of 256, Adam optimization, mean-squared-error loss, and learning rates of 10⁻³ during fusion-head training and 10^-4^ and 3 × 10^-4^ for encoder and fusion-head fine-tuning, respectively. To assess whether protein-level and site-level representations combine approximately linearly or require additional nonlinear integration, we compared the full nonlinear fusion model with a linear additive fusion model in which the nonlinear fusion head was replaced with a single linear layer operating on the concatenated encoder outputs.

## Supplementary Figures

**Figure S1.**
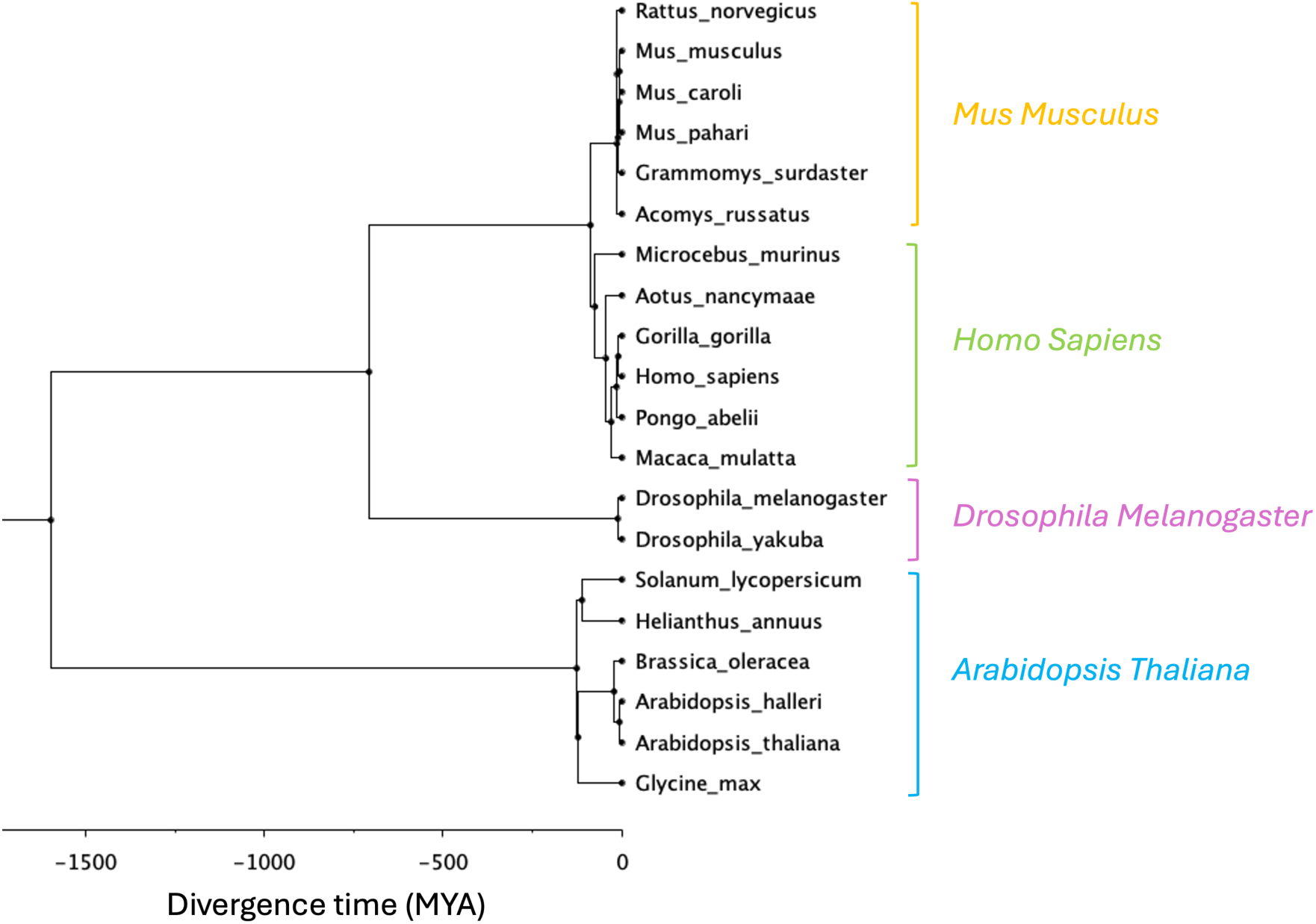
Phylogenetic relationships and divergence times of the species used to estimate protein evolutionary rates. For each focal species—*Homo sapiens*, *Mus musculus*, *Drosophila melanogaster*, and *Arabidopsis thaliana*—we selected a set of closely and progressively diverging secondary species (shown as grouped clades) to estimate substitution rates. Divergence times (in million years ago, MYA) are indicated along the X-axis.

**Figure S2:**
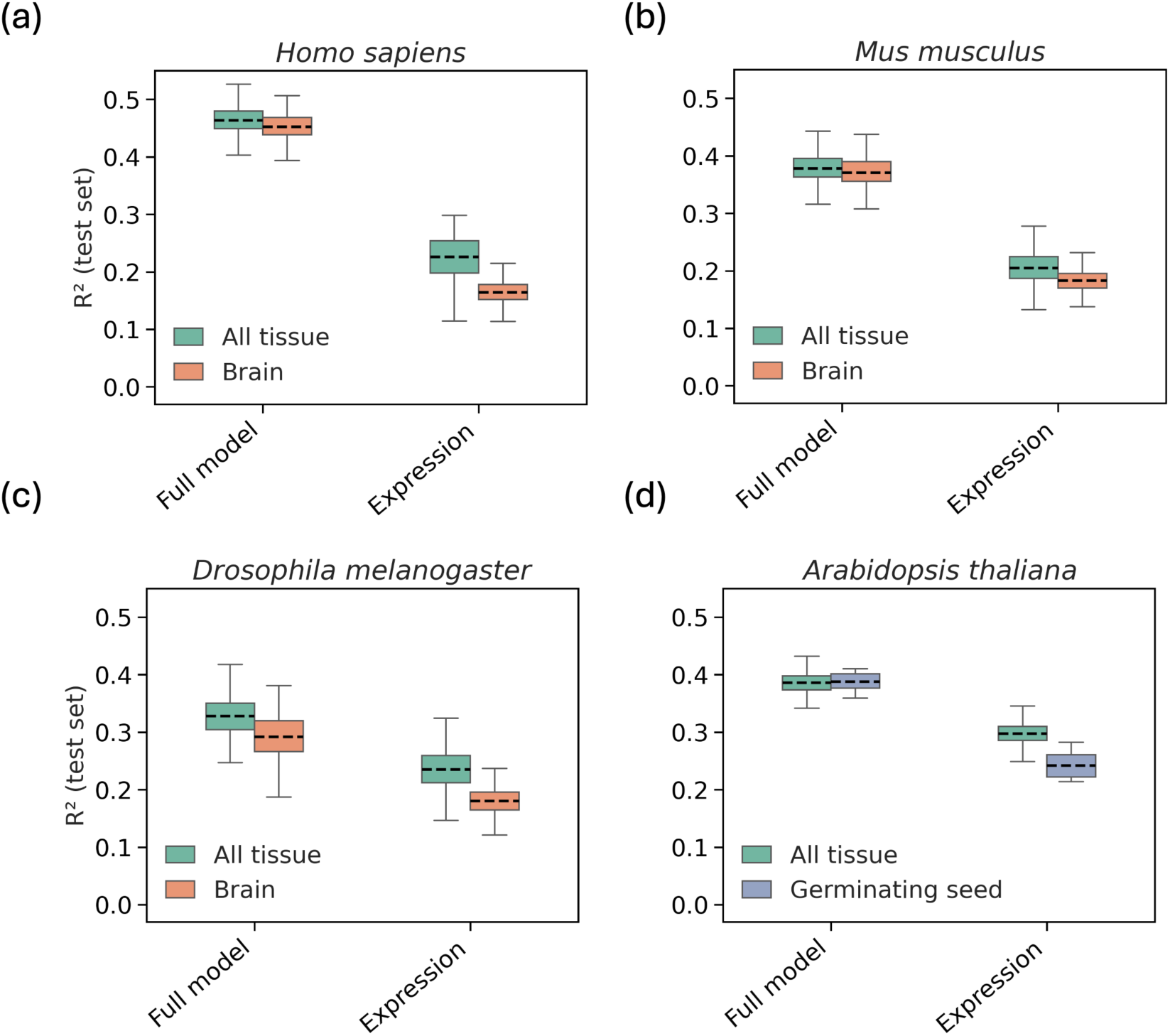
Model performance comparison between using expression across all tissues versus the single dominant tissue driving the expression–evolutionary rate (E–R) relationship. Predictive performance (test-set 𝑅^2^) of the full model and expression-only models across representative eukaryotic lineages: *Homo sapiens*, *Mus musculus*, *Drosophila melanogaster*, and *Arabidopsis thaliana*. For mammals and *Drosophila melanogaster*, the single-tissue model used brain expression, whereas for *Arabidopsis thaliana* it used germinating seed expression. Empirical evolutionary rates for each species were estimated using a star phylogeny (see Methods). Points summarize the distribution of 𝑅^2^values across 500 independent runs with random train/validation/test gene splits; boxes indicate the mean and standard deviation.

**Figure S3.**
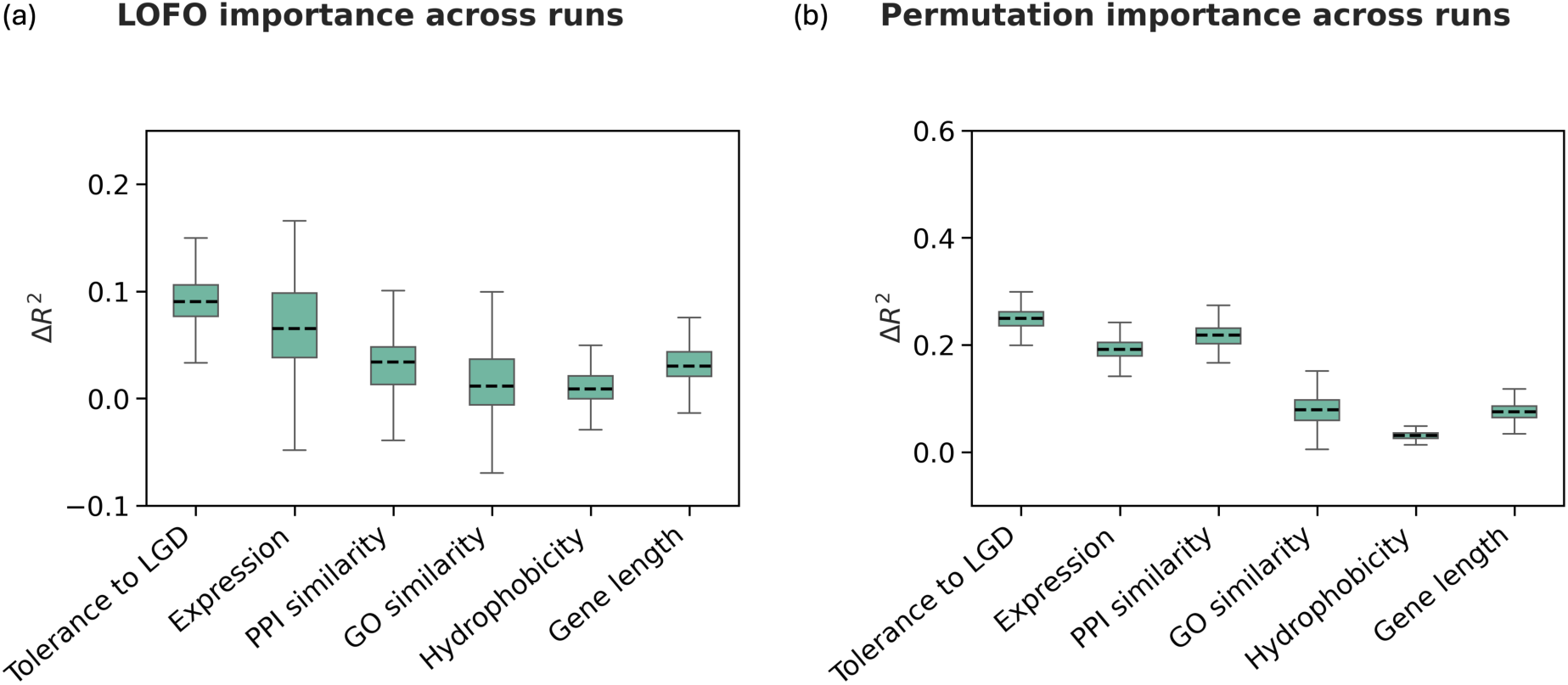
Feature importance across runs based on LOFO and permutation analyses. Each panel shows the contribution of different feature blocks—gene dosage (tolerance to LGD mutations), expression, PPI network context, GO-based function, hydrophobicity, and gene length to model performance, quantified as the change in 𝑅^2^ (Δ𝑅^2^). (a) LOFO (leave-one-feature-out): each feature block is removed, the model is retrained, and the resulting decrease in 𝑅^2^is used to estimate its contribution. (b) Permutation importance: each feature block is randomized, and the decrease in 𝑅^2^reflects how much the model relies on that feature for prediction. Both analyses were repeated over 500 runs, where genes were randomly split into training, validation, and test sets in each run. Boxplots show the distribution of Δ𝑅^2^across runs. The mean and standard deviations are shown.

**Figure S4:**
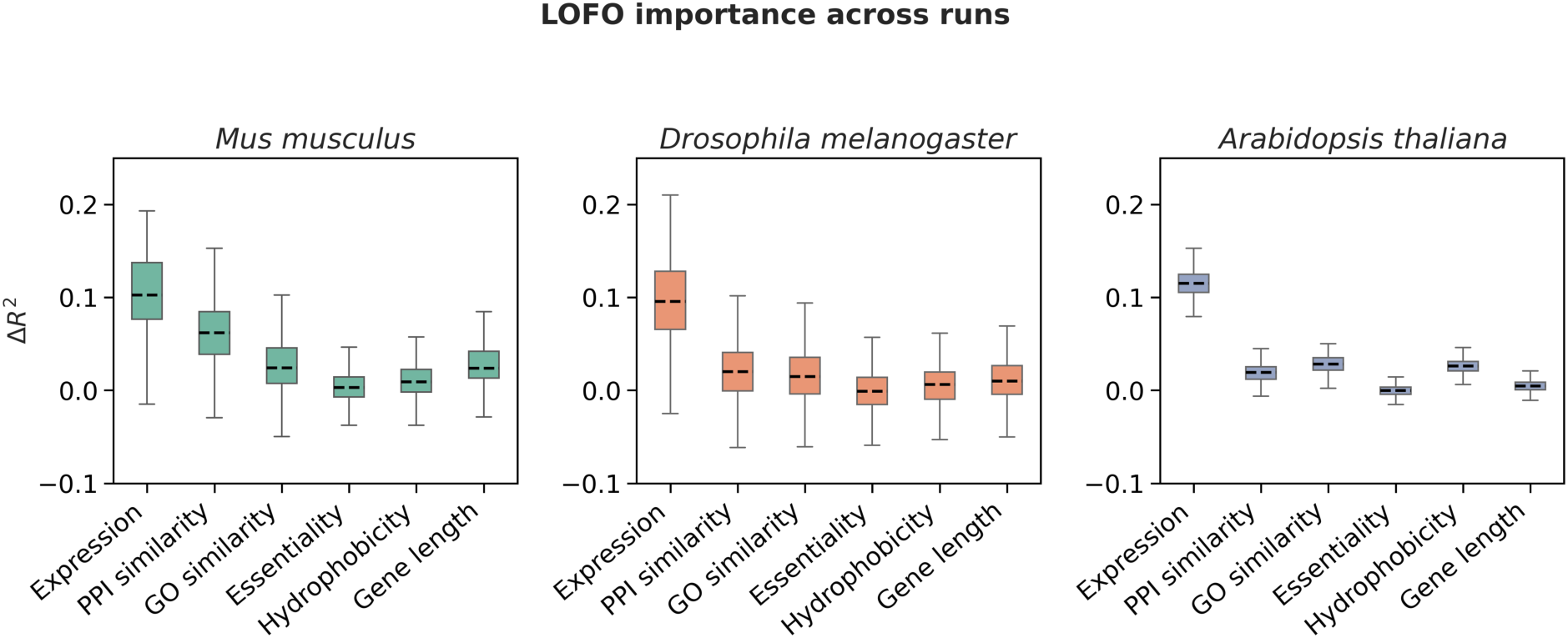
The individual feature importance in the model prediction across different species using LOFO (leaving one feature out) analysis. We estimate the contribution of each feature block (Expression, PPI, GO, Essentiality, Hydrophobicity and Gene length) by dropping the feature and retraining the network, and calculating the drop in 𝑅^2^. The analysis is shown for *Mus musculus*, *Drosophila melanogaster* and *Arabidopsis thaliana*. The method is repeated across 500 runs in which, within each run, genes are randomly split into training/validation/test sets. The mean and standard deviations are shown.

**Figure S5:**
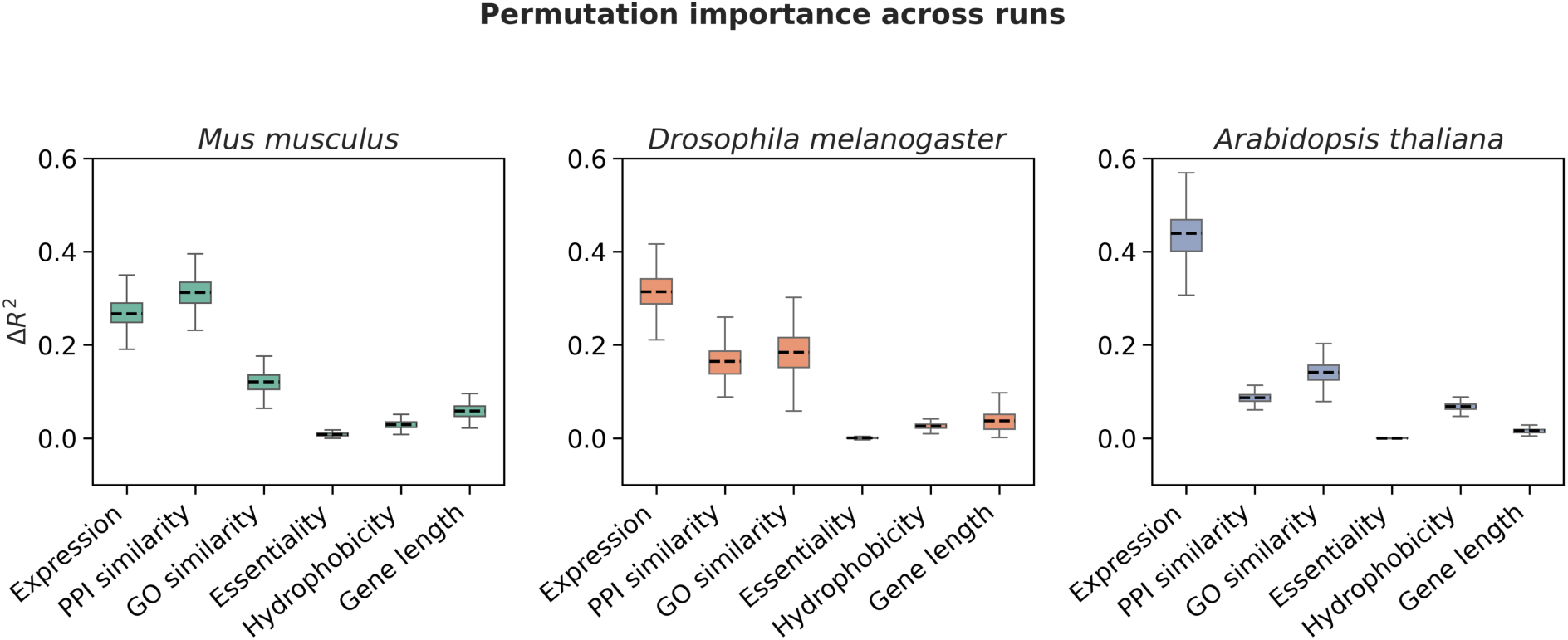
The individual feature importance in the model prediction across different species using permutation analysis. We estimate the contribution of each feature block (Expression, PPI, GO, Essentiality, Hydrophobicity and Gene length) by randomizing the feature block and calculating the drop in 𝑅^2^, which reflects how much the model uses that feature for prediction. The analysis is shown for *Mus musculus*, *Drosophila melanogaster* and *Arabidopsis thaliana*. The method is repeated across 500 runs in which, within each run, genes are randomly split into training/validation/test sets. The mean and standard deviations are shown.

**Figure S6:**
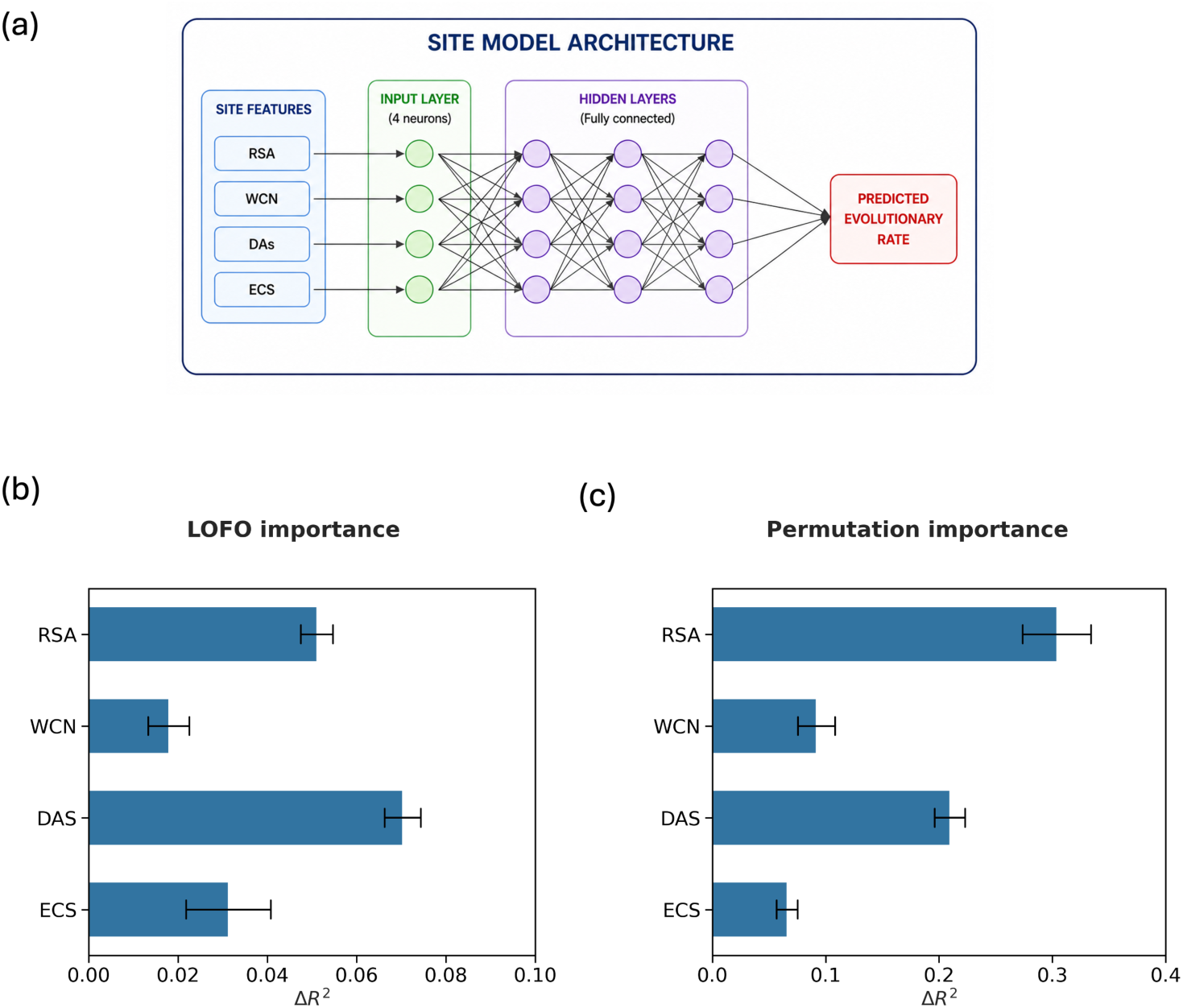
Feature importance in the prediction of relative evolutionary rates across sites. (a) Schematic of the site-level neural network used to predict the relative evolutionary rate of residues within a protein. Site features (RSA, WCN, DAS, ECS) are provided as input and passed through fully connected hidden layers to predict the evolutionary rate at each site. (b) Feature importance estimated using leave-one-feature-out (LOFO) analysis, where each feature is removed from the model and the resulting change in 𝑅^2^ is measured. (c) Feature importance estimated using permutation analysis, where each feature is randomly shuffled in the test set and the resulting change in 𝑅^2^ is measured.

**Table S1:** Model performance and feature importance across species. Model performance is reported as the mean test-set R² ± standard deviation across 500 runs for each feature-specific model. Feature importance is quantified using both permutation importance and leave-one-feature-out (LOFO) analysis, reported as the change in predictive performance (ΔR²) upon perturbation or removal of each feature. Dashes indicate values not applicable for the full model. In Homo sapiens, gene constraint is represented by tolerance to loss-of-function mutations (Tolerance (LGD)), whereas in other species it is represented by essentiality.

| Species | Feature | Model Performance | LOFO Importance | Permutation Importance |
| --- | --- | --- | --- | --- |
| Homo sapiens | Full model | $0.463 \pm 0.024$ | - | - |
| Homo sapiens | Tolerance (LGD) | 0.251 ± 0.020 | 0.090 ± 0.041 | 0.249 ± 0.020 |
| Homo sapiens | Expression | 0.226 ± 0.039 | 0.065 ± 0.044 | 0.191 ± 0.021 |
| Homo sapiens | PPI | 0.211 ± 0.025 | 0.034 ± 0.052 | 0.218 ± 0.022 |
| Homo sapiens | GO | 0.130 ± 0.021 | 0.012 ± 0.045 | 0.079 ± 0.025 |
| Homo sapiens | Hydrophobicity | 0.005 ± 0.003 | 0.009 ± 0.041 | 0.031 ± 0.007 |
| Homo sapiens | Gene length | 0.005 ± 0.004 | 0.030 ± 0.039 | 0.075 ± 0.015 |
| Mus musculus | Full model | 0.378 ± 0.025 | - | - |
| Mus musculus | Essentiality | 0.023 ± 0.008 | 0.003 ± 0.049 | 0.008 ± 0.004 |
| Mus musculus | Expression | 0.204 ± 0.028 | 0.102 ± 0.045 | 0.267 ± 0.036 |
| Mus musculus | PPI | 0.207 ± 0.024 | 0.062 ± 0.051 | 0.312 ± 0.033 |
| Mus musculus | GO | 0.109 ± 0.021 | 0.024 ± 0.048 | 0.121 ± 0.023 |
| Mus musculus | Hydrophobicity | 0.002 ± 0.002 | 0.009 ± 0.045 | 0.029 ± 0.008 |
| Mus musculus | Gene length | 0.009 ± 0.004 | 0.024 ± 0.045 | 0.058 ± 0.016 |
| Drosophila melanogaster | Full model | 0.328 ± 0.033 | - | - |
| Drosophila melanogaster | Essentiality | 0.071 ± 0.014 | -0.001 ± 0.041 | 0.000 ± 0.001 |
| Drosophila melanogaster | Expression | 0.235 ± 0.034 | 0.095 ± 0.045 | 0.314 ± 0.043 |
| Drosophila melanogaster | PPI | 0.176 ± 0.033 | 0.020 ± 0.038 | 0.165 ± 0.036 |
| Drosophila melanogaster | GO | 0.149 ± 0.027 | 0.015 ± 0.036 | 0.184 ± 0.049 |
| Drosophila melanogaster | Hydrophobicity | 0.009 ± 0.005 | 0.006 ± 0.037 | 0.026 ± 0.006 |
| Drosophila melanogaster | Gene length | 0.002 ± 0.002 | 0.010 ± 0.035 | 0.037 ± 0.020 |
| Arabidopsis thaliana | Full model | 0.386 ± 0.018 | - | - |
| Arabidopsis thaliana | Essentiality | 0.004 ± 0.003 | -0.000 ± 0.006 | -0.000 ± 0.000 |
| Arabidopsis thaliana | Expression | 0.297 ± 0.019 | 0.115 ± 0.015 | 0.438 ± 0.053 |
| Arabidopsis thaliana | PPI | 0.132 ± 0.023 | 0.019 ± 0.011 | 0.087 ± 0.010 |
| Arabidopsis thaliana | GO | 0.196 ± 0.017 | 0.028 ± 0.010 | 0.141 ± 0.024 |
| Arabidopsis thaliana | Hydrophobicity | 0.027 ± 0.007 | 0.026 ± 0.008 | 0.067 ± 0.008 |
| Arabidopsis thaliana | Gene length | 0.001 ± 0.001 | 0.005 ± 0.007 | 0.016 ± 0.006 |

**Table S2:**
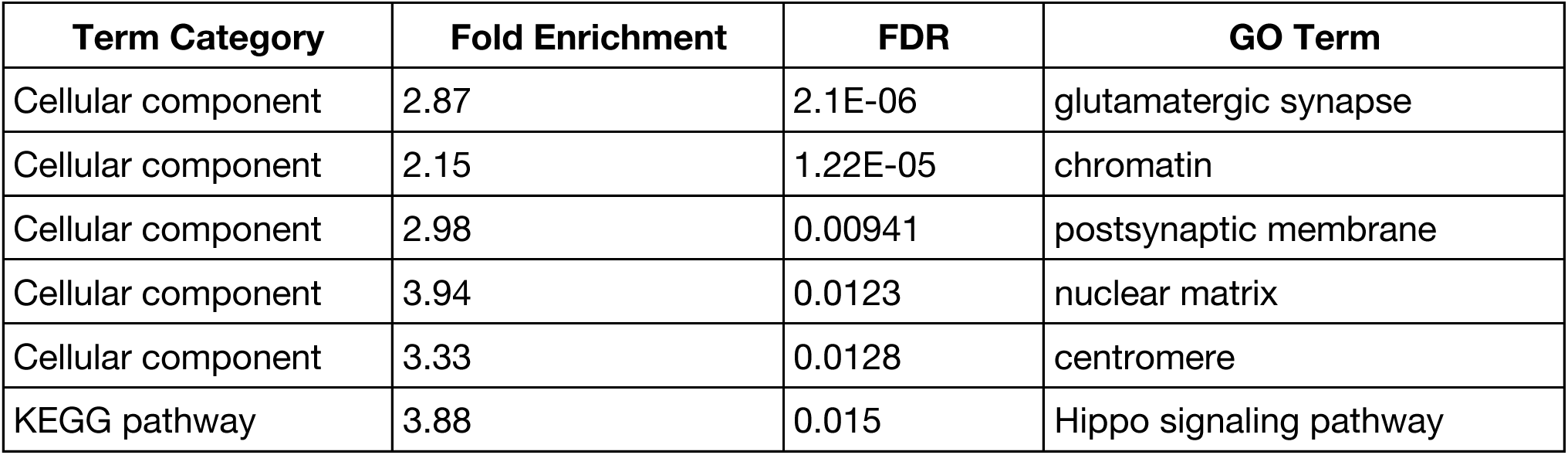
GO and pathway enrichment analysis using DAVID for the top 500 overpredicted proteins. The table shows enriched categories including GO biological processes, molecular functions, cellular components, and KEGG pathways, along with their corresponding terms, fold enrichment, and FDR values. Proteins were ranked based on the average difference between predicted and empirical evolutionary rates across runs, where the model overestimates the rate. Enriched terms highlight biological processes and cellular components associated with proteins whose evolutionary rates are systematically overestimated by the model.

**Table S3:** GO and pathway enrichment analysis using DAVID for the top 500 underpredicted proteins. The table shows enriched categories including GO biological processes, molecular functions, cellular components, and KEGG pathways, along with their corresponding terms, fold enrichment, and FDR values. Proteins were ranked based on the average difference between predicted and empirical evolutionary rates across runs, where the model underestimates the rate. Enriched terms highlight biological processes and molecular functions associated with proteins whose evolutionary rates are systematically underestimated by the model.

| Term Category | Fold Enrichment | FDR | GO Term |
| --- | --- | --- | --- |
| Molecular function | 3.18 | 5.28E-10 | sequence-specific double-stranded DNA binding |
| Cellular component | 2.28 | 4.88E-07 | chromatin |
| Molecular function | 3.83 | 2.39E-06 | transcription cis-regulatory region binding |
| Biological process | 2.24 | 3.16E-06 | positive regulation of transcription by RNA polymerase II |
| Molecular function | 2.49 | 2.11E-05 | Repressor |
| Molecular function | 2.36 | 4.07E-05 | DNA-binding transcription factor activity |
| Molecular function | 2.07 | 4.07E-05 | RNA polymerase II cis-regulatory region sequence-specific DNA binding |
| Molecular function | 1.85 | 4.07E-05 | RNA binding |
| Biological process | 1.97 | 7.28E-05 | regulation of DNA-templated transcription |
| Molecular function | 2.24 | 7.13E-05 | Activator |
| Biological process | 2.16 | 0.000192 | negative regulation of transcription by RNA polymerase II |
| Molecular function | 2.67 | 0.000265 | DNA-binding transcription activator activity, RNA polymerase II-specific |
| Biological process | 1.85 | 0.000287 | regulation of transcription by RNA polymerase II |
| Molecular function | 2.09 | 0.000356 | RNA-binding |
| Biological process | 2.24 | 0.00114 | positive regulation of DNA-templated transcription |
| Molecular function | 1.82 | 0.00132 | DNA-binding transcription factor activity, RNA polymerase II-specific |
| Biological process | 2.02 | 0.00165 | Host-virus interaction |
| Molecular function | 2.91 | 0.00264 | DNA-binding transcription repressor activity, RNA polymerase II-specific |
| Molecular function | 2.31 | 0.00482 | protein kinase binding |
| Biological process | 3.26 | 0.00583 | protein stabilization |
| Cellular component | 2.38 | 0.0137 | nuclear speck |
| Molecular function | 1.65 | 0.0171 | Developmental protein |

## Acknowledgements

This work was supported by grant R35GM131884 from the National Institutes of Health.

